# FastHer: fast and accurate estimator of local heritability using summary statistics

**DOI:** 10.64898/2025.12.24.696369

**Authors:** Gulnara R. Svishcheva, Yakov A. Tsepilov, Tatiana I. Axenovich

## Abstract

We present FastHer, a method for estimating local heritability using GWAS summary statistics and linkage disequilibrium (LD) matrices. FastHer performs exact restricted maximum-likelihood estimation matching the accuracy of individual-level methods. Analytical transformations ensure high computational efficiency. FastHer incorporates SNP weights that define the contribution of each SNP to heritability. The introduction of a model-data concordance parameter enables the adaptation of reference LD matrices when in-sample LD matrices are unavailable.

## Main

Local heritability (*h*^2^) quantifies the proportion of trait variance attributable to a genomic region. Restricted maximum likelihood (REML), implemented as GREML, is a widely used reference method, yielding precise *h*^2^ estimates from individual-level data^1^. Limited availability of such data motivates the development of *h*^2^ estimators that use linkage disequilibrium (LD) information and GWAS summary statistics (*z*-statistics, minor allele frequencies (MAF) and sample sizes).

Summary-statistics methods differ in how LD information is represented (as LD scores or as an LD correlation matrix), and in the estimation approach (method of moments or maximum likelihood). The method of moments, as implemented in LDSC, GRE, HESS, EHE and RHE-reg^2-6^, generally yields less precise heritability estimates^7^ than REML. Among maximum-likelihood methods, HEELS^8^, implementing exact REML, achieves accuracy comparable to GREML. Nevertheless, HEELS is computationally intensive. Another method, LDAK-SumHer^9^ (hereafter SumHer), is an LD-score-based method and maximizes an approximate likelihood. SumHer offers a key advantage over other summary-statistics methods: it allows the user to specify the heritability model, defining the assumed genetic architecture.

We present FastHer, a method that uses *z*-statistics and a precomputed LD matrix (Pearson’s SNP-SNP correlations), requiring no raw genotype data. FastHer performs exact REML, achieving accuracy comparable to GREML and HEELS. Like SumHer, FastHer allows the user to specify SNP weights that define how heritability is distributed across SNPs, and it supports reference LD matrices. FastHer’s high computational efficiency stems from analytical transformations:

∘ The individual-level Gaussian log-likelihood is reformulated in terms of summary statistics by decomposing it into two orthogonal components: one depends solely on the residual variance 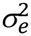, while the other depends on 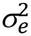 and the genetic variance 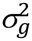.
∘ For any fixed 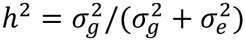, the total model-implied variance 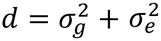 has an analytical solution. Profiling out *d* in this way enables exact REML estimation of (*h*^2^, *d*) via a one-parameter search over *h*^2^.
∘ An eigen-decomposition of the LD matrix, performed once at cost *O*(*m*^3^), reduces the cost of each subsequent iteration to *O*(*r*), where *r* is the effective rank of the (*m*×*m*) LD matrix.

The parameter *d* reflects the model-data concordance: *d*=1 indicates the equality of the model-implied and sample phenotypic variances. Furthermore, 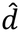 guides an adaptive eigenvalue truncation which regularizes reference LD matrices when in-sample matrices are unavailable: the components of the reference LD matrix with the smallest eigenvalues are removed one-by-one until 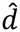 is close to 1 (Wald statistic ≤1.96; two-sided *α*=0.05).

Details are provided in *Methods* and Supplementary Note 1 (Supplementary Table 2).

We compared FastHer with HEELS and SumHer from a methodological perspective (Supplementary Table 1). In all methods, GWAS *z*-statistics serve as input. FastHer and HEELS directly use a precomputed LD matrix, whereas SumHer requires precomputed, model-specific tagging scores. Both FastHer and HEELS perform exact REML, with a per-iteration cost *O*(*r*) and *O*(*m*^3^), respectively. Both estimate standard errors (SE) analytically. In contrast, SumHer maximizes an approximate likelihood based on a gamma distribution of the squared *z*-statistics, and SE(*h*^2^) is estimated empirically. SumHer is designed for common SNPs (MAF>0.005), whereas FastHer imposes no MAF restriction. Similar to SumHer (but unlike HEELS), FastHer supports user-specified SNP weights and includes an eigenvalue-truncation procedure for regularizing reference LD matrices when in-sample matrices are unavailable.

To compare FastHer and HEELS numerically, we considered phenotypes simulated using genotypes from the 1000 Genomes Project (1000G)^10^ and real UK Biobank (UKB) GWAS summary statistics with in-sample LD matrices for five binary cardiometabolic traits and three continuous biomarkers (Supplementary Notes 2–3). In both cases, region-level estimates of *h*^2^ and SE(*h*^2^) agreed closely between the two methods (Fig. 1a,b; Supplementary Table 3; Supplementary Figs 1–7). We compared the runtime of FastHer and HEELS in simulations: the advantage of FastHer increased with the number of SNPs, and for the largest gene (10,968 SNPs; *h*^2^=0.025), FastHer was ∼100-fold faster than HEELS (median 6.4 min vs 10.7 h; Fig. 1c; Supplementary Note 4, Supplementary Fig. 8).

**Figure 1.**
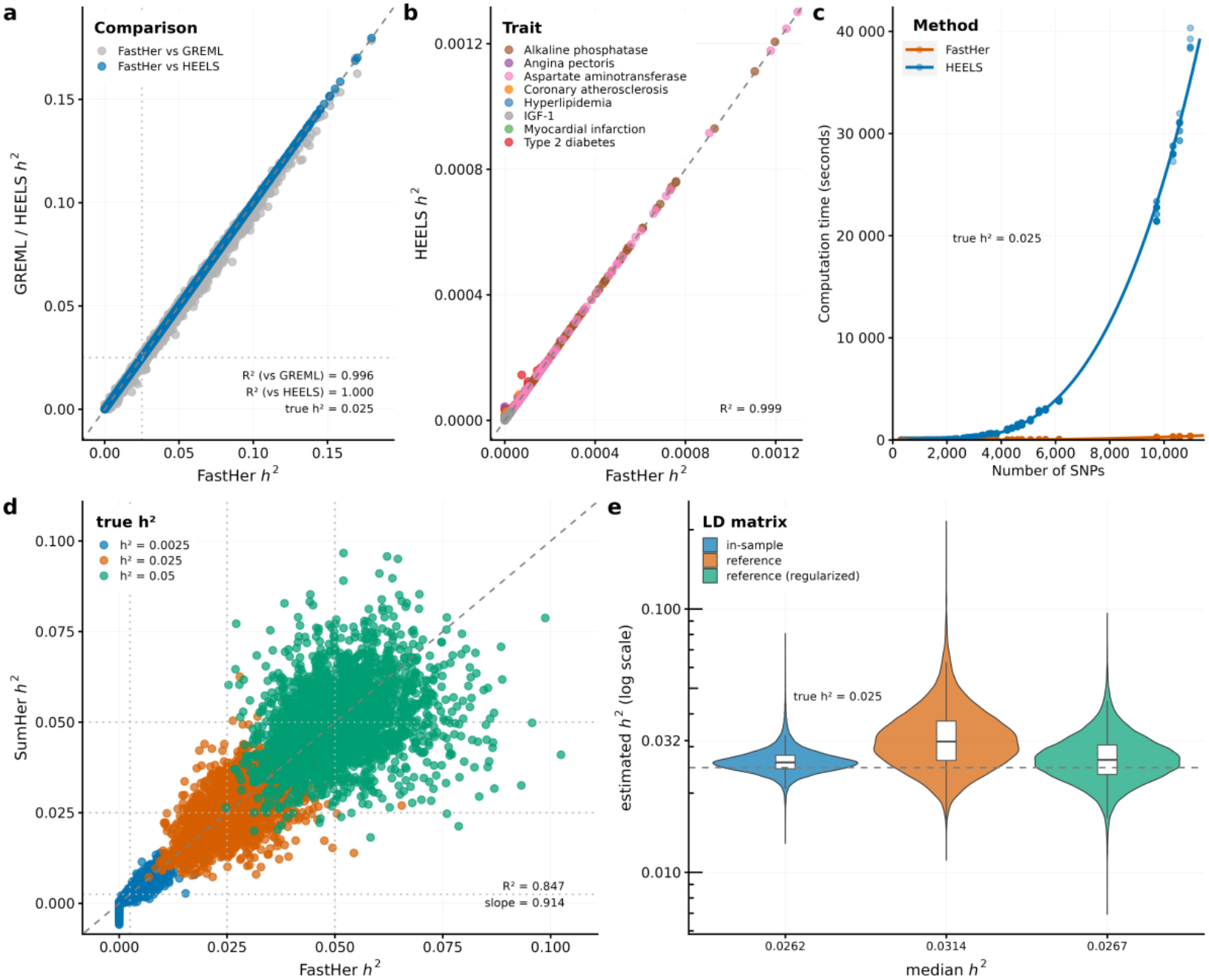
Benchmarking FastHer and reference LD regularization. **a**, *h*^2^ from FastHer versus GREML/HEELS (simulated phenotypes, 1000G, GCTA model). **b**, *h*^2^ from FastHer versus HEELS (UKB summary statistics; the Wald test did not reject *d*=1 for all pairs). **c**, Runtime of FastHer and HEELS versus number of SNPs (*h*^2^=0.025; simulated summary statistics, UKB LD matrices, GCTA model). **d**, *h*^2^ from FastHer versus SumHer (simulated phenotypes, 1000G, LDAK model). **e**, Distribution of *h*^2^ obtained with in-sample, unregularized and regularized reference LD matrices.

To compare FastHer and SumHer, we simulated phenotypes assuming unequal SNP contributions to heritability (*Methods*). Both methods yielded similar estimates for *h*^2^ and SE(*h*^2^), with Pearson correlation coefficients of 0.92 and 0.94, respectively (Fig. 1d; Supplementary Note 5, Supplementary Fig. 9; Supplementary Table 4).

We also evaluated regularization of the reference LD matrices. Across loci, we simulated summary statistics from the UKB LD matrices and applied FastHer using these matrices and the 1000G reference matrices, with and without regularization (*Methods*). Without regularization, *h*^2^ estimates were substantially inflated, whereas with regularization they were close to those obtained using the original matrices (Fig. 1e; Supplementary Note 6, Supplementary Fig. 10).

Thus, to our knowledge, FastHer is the only method that combines exact REML from GWAS summary statistics and LD correlation matrices, SNP weighting, high computational efficiency for large loci and regularization of reference LD matrices. Future extensions could encompass alternative heritability models and more complex genetic architectures.

## Methods

### Individual-level model specification

Let *y* ∈ ℝ^*n*^ be a vector of phenotypes at *n* individuals, adjusted for covariates and standardized, *y*^*T*^*y* = *n*. For a given locus spanning *m* SNPs, we assume that *y* follows a Gaussian distribution

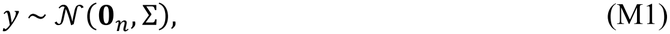

where the local covariance matrix Σ is

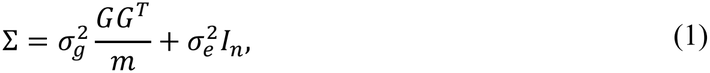

*G* ∈ ℝ^*n*×*m*^ is a standardized genotype matrix, with *r* = rank(*G*) ≤ min(*n*– 1, *m*); 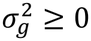 is the genetic variance of *y*; 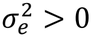 is the residual variance of *y*; and *I* _*n*_ ∈ ℝ^*n*×*n*^ is the identity matrix.

Model (M1) can equivalently be parametrized by the local heritability *h*^2^ and a scaling parameter *d*: 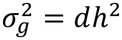 and 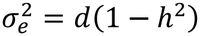, defining a one-to-one mapping from 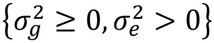 onto {*h*^2^ ∈ [0,1), *d* > 0}. Then 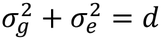, and Σ = *d*Σ_0_ with 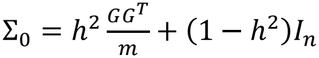. Here *d* is the ratio of the model-implied marginal variance of *y* to its observed variance, which equals 1. For fixed *h*^2^, the REML estimate of *d*, which is also the method-of-moments estimate, is

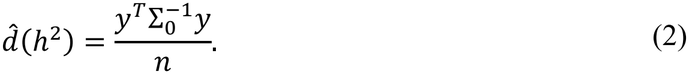

*d* serves as a concordance parameter: under a correctly specified model, 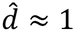, whereas it can deviate substantially from 1 when the model is misspecified (e.g., due to population stratification, genotyping errors, or mismatched reference LD matrices).

For *y*, the −2 log-likelihood is

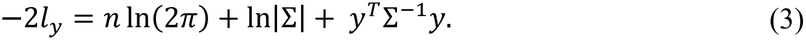

### Orthogonal decomposition of log-likelihood

We introduce the orthogonal projection matrix onto the null space of *G*^*T*^:

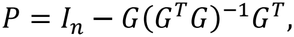

where (·)^−1^ is the Moore–Penrose pseudoinverse. *P* is idempotent (*P*^2^ = *P*), symmetric (*P*^*T*^=*P*), satisfies *PG*=**0**, and rank(*P*) = *n*– *r* ≥ 1. It splits the phenotype vector into orthogonal vectors:

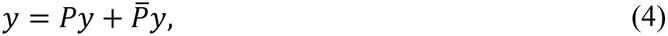

where 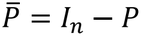; note that *Py* lies in the (*n−r*)-dimensional subspace orthogonal to the columns of *G*, while 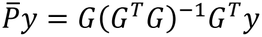 lies in the *r*-dimensional column space of *G*.

#### Lemma 1.

Under model (M1), (*i*) 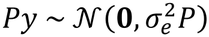, a degenerate Gaussian distribution in the (*n−r*)-dimensional subspace orthogonal to the columns of *G*, whose distribution depends only on 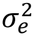; (*ii*) *Py* and 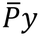 are independent.

*Proof*. (*i*) *Py* is a linear transformation of a Gaussian vector, and 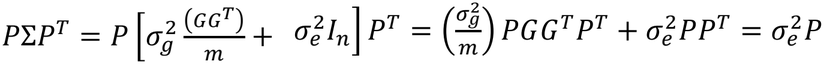, since *PG*=**0** and *PP*^*T*^ = *P*. (*ii*) 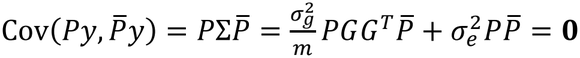, since *PG=***0** and 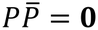. The vectors are jointly Gaussian, hence uncorrelated implies independent. ▪

By **Lemma 1**, the −2 log-likelihood (3) of *y* is the sum of the −2 log-likelihoods of the two independent components from (4),

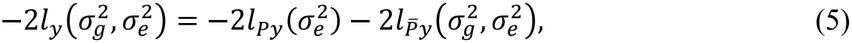

where, since rank(*P*) = *n* – *r*,

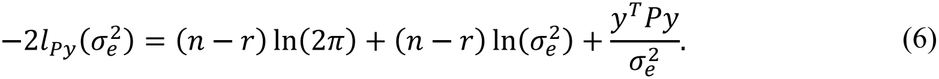

In (5), the first component depends only on 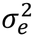, whereas the second depends on 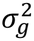 and 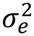; both components are expressed exactly in terms of summary statistics in the next section.

### Log-likelihood in terms of summary statistics

Define the marginal *z*-statistics and the LD matrix as

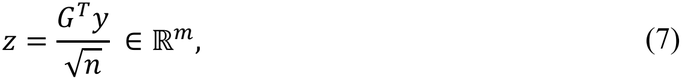

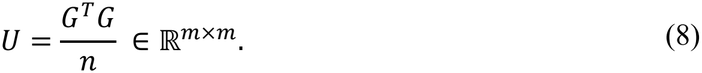

*U* is symmetric positive-semidefinite with rank *r* ≤ min(*n* − 1, *m*); *U*^−*l*^ denotes its Moore–Penrose pseudoinverse raised to the power *l*>0. Under model (M1), 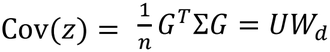, so *z* ∼ *N*(0_*m*_, *UW*_*d*_), where

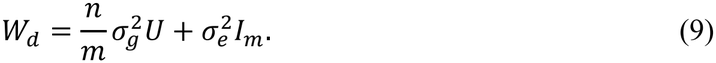

As 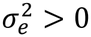, *W*_*d*_ is symmetric positive-definite, so 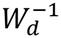 and ln|*W*_*d*_ | are always well-defined; if *r<m*, the distribution of *z* is degenerate, since rank(*UW*_*d*_) = *r* < *m* . Since *W*_*d*_ (9) is a polynomial in *U, W*_*d*_ commutes with *U*, as well as with its powers.

We introduce *z*_*u*_ = *U*^−1/2^*z* as the vector of LD-adjusted *z*-statistics, so that 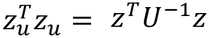 and 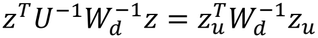. Then, according to (7) and (8),

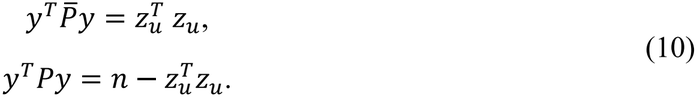

#### Theorem 1 (exact decomposition)

Define *F*_1_ and *F*_2_ as the −2 log-likelihoods of 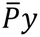 and *Py*, respectively:

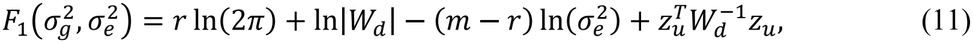

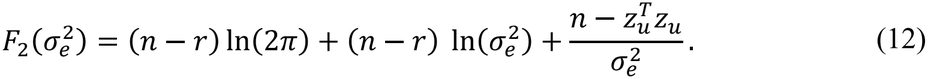

Then

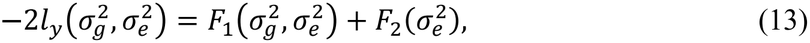

where *F*_1_ depends on both parameters through *W*_*d*_, and *F*_2_ depends only on 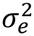. The decomposition (13) is exact.

*Proof*. By (6) and (10), −2*l*_*Py*_ = *F*_2_. It remains to show that −2*l*_*y*_ = *F*_1_ + *F*_2_; then 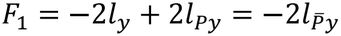 by (5). Applying the matrix determinant lemma to Σ (1) gives 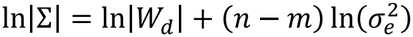, and the Woodbury identity gives 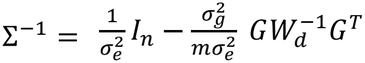. Therefore, using *y*^*T*^*y* = *n*,

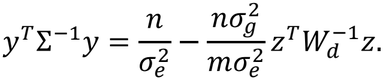

From (9), 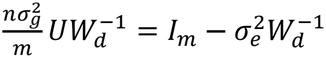; since *z* lies in the column space of *G, z* = *UU*^−1^*z*, and

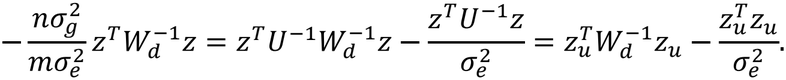

Substituting,

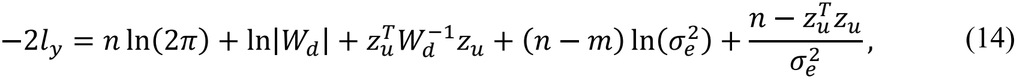

which equals *F*_1_ + *F*_2_. ▪

We estimate *d* in terms of summary statistics. Since 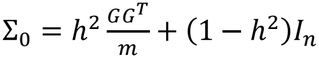 and 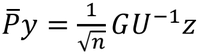, the same argument gives 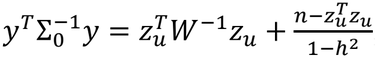, so that (2) becomes

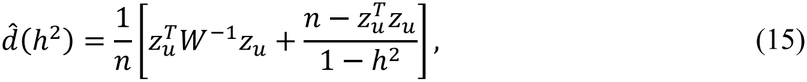

where 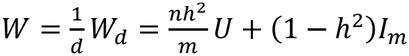.

### Exact *d*-profiling: *h*^2^ estimation

Substituting 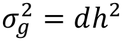 and 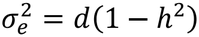 into (14) gives

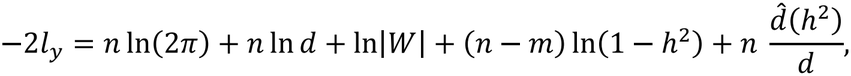

where *W* and 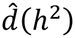 are defined in (15). Because *d* has a closed-form expression in terms of *h*^2^ – its value at the minimum of −2*l*_*y*_ = *F*_1_ + *F*_2_ over *d*, for fixed *h*^2^, namely 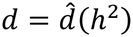 – substituting 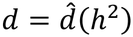 back eliminates *d* entirely, leaving a function of *h*^2^ alone: the exact profile −2 log-likelihood (up to an additive constant),

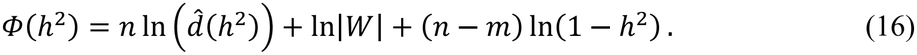

The profile-REML estimate is 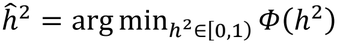, with 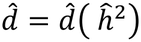. Since the parametrizations 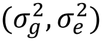and (*h*^2^, *d*) are one-to-one, and the minimum over *d* is attained in closed form at 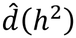 (provided 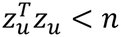, the profile-REML estimate coincides exactly with the two-parameter REML estimate.

### Relationship to HESS and GRE

Minimizing *F*_2_ (12) over 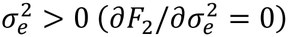 gives, provided 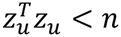,

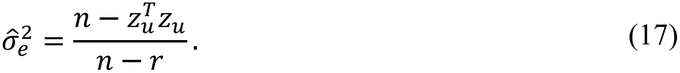

The second derivative, 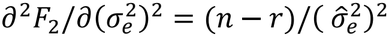, is positive at 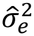, confirming a minimum. Setting *d*=1 (full model-data concordance) in 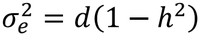 gives 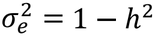, and (17) yields

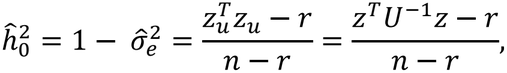

which coincides with the HESS and GRE estimators^3,5^. Notably, this estimator is invariant to assumptions about the joint distribution of SNP effects (including SNP weighting), since it measures the deviation of the Mahalanobis norm *z*^*T*^*U*^−1^*z* from its null expectation *r* (at *h*^2^=0). Thus, these estimators correspond to minimizing *F*_2_ alone with *d* fixed at 1: they depend on the data only through the scalar *z*^*T*^*U*^−1^*z*, whereas the REML estimator additionally uses *F*_1_ and estimates *d*.

### Computational efficiency

Let 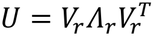 be the eigen-decomposition of *U*, where *Λ*_*r*_ = diag(*λ*_1_, …, *λ*_*r*_) contains the *r* strictly positive eigenvalues and *V*_*r*_ = (*v*_1_, …, *v*_*r*_) ∈ ℝ^*m*×*r*^ is the matrix of the corresponding eigenvectors 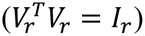. It is computed once per locus at *O*(*m*^3^) cost. The coordinates of *z*_*u*_ in this eigenbasis,

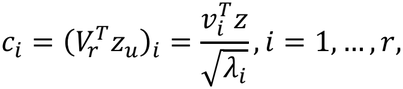

are computed once at *O*(*mr*) cost, and 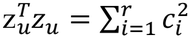.

In this eigenbasis, *F*_1_ has a simple form. The eigenvalues of *W* are 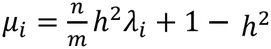 for *i*=1,…,*r*, and 1 − *h*^2^ with multiplicity *m−r*. Therefore

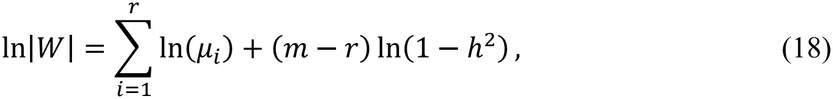

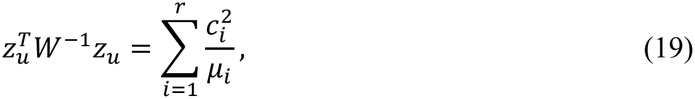

and Eq. (11) becomes

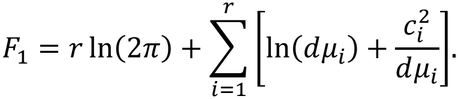

This is the −2 log-likelihood of *r* independent variables *c*_*i*_∼*N*(0, *dμ*_*i*_): by (9),

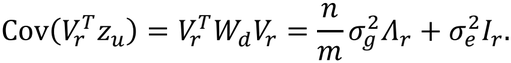

Substituting (18) and (19) into (15) and (16) gives 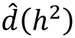 and the profile objective in the eigenbasis,

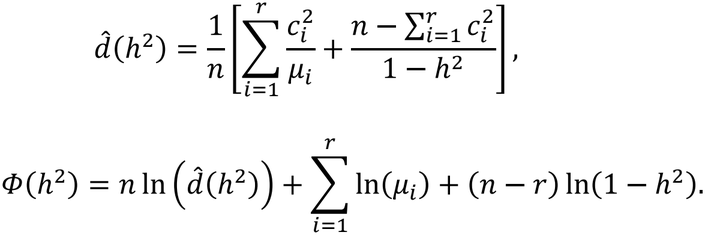

Each evaluation of *Φ*(*h*^2^) requires *O*(*r*) operations. The estimation thus reduces to a one-dimensional search over *h*^2^ with *O*(*r*) cost per evaluation, once *U* has been decomposed.

### Model with SNP weighting

Following the LDAK framework^11^ and the SumHer approach for summary statistics^12^, we allow the per-SNP heritability to depend on MAF, which improves *h*^2^ estimation under non-uniform genetic architectures^13^. In the LDAK model with *--power −0*.*25*, the expected heritability of SNP *i* is proportional to [*MAF*_*i*_(1 − *MAF*_*i*_)]^0.75^. To incorporate this assumption into our framework, we define the weight for SNP *i* as

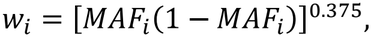

so that 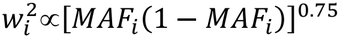. The weights are normalized so that 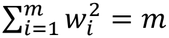 i.e. 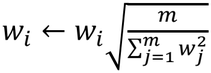 . Thus the mean of 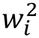 equals 1 and *h*^2^ keeps its scale. Let *D* = diag(*w*_1_, …, *w*_*m*_). The weighted model is the original model with the genotype matrix *G* replaced by *GD*, so that *U* and *z* are replaced by

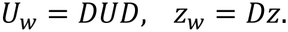

The eigen-decomposition of *U*_*w*_ is computed once, so computational efficiency is preserved. MAF values are taken from the same summary statistics as *z*.

### Regularization for reference LD matrices

When a reference LD matrix (e.g., 1000G) replaces the in-sample matrix, estimates may be biased by population mismatch and sampling noise. In practice, the true *h*^2^ is unknown, so the accuracy of *h*^2^ cannot be assessed directly. The second model parameter, *d*, is expected to equal 1 under model-data concordance. FastHer therefore tests the deviation of 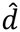 from 1 with a Wald statistic, 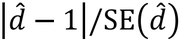, at a two-sided *α*=0.05 (critical value 1.96).

Let *U*_*ref*_ = *VΛV*^*T*^ be the eigen-decomposition of the reference matrix, with eigenvalues *λ*_1_ ≥ ⋯ ≥ *λ*_*m*_; negative eigenvalues are set to zero. For a threshold *τ*, the retained rank *r* is the number of leading components whose cumulative share of the total LD variance is below *τ*:

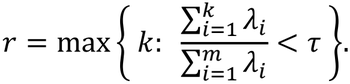

*U*_*ref*_ is then replaced by an eigenvalue-truncated matrix *U*_*r*_ = *V*_*r*_*Λ*_*r*_*V*^*T*^. This procedure removes the components with the smallest eigenvalues, which are the most sensitive to sampling noise: they enter any quadratic form involving 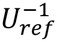 with weights 1/*λ*_*i*_, so underestimated eigenvalues inflate such forms and distort 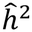 and 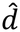.

We start with *τ*=0.97 and compute 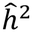, 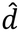) with the current *U*_*r*_. If the Wald statistic exceeds 1.96, *r* is reduced by one component and the estimation is repeated. The procedure stops when the Wald statistic is ≤1.96, or when the retained fraction falls to 0.95, or when *r* reaches 6. 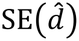 is obtained from the Fisher information matrix (Supplementary Note 1). Truncation reduces the bias caused by a mismatched reference at the cost of precision (Fig. 1e).

In simulations, we also computed 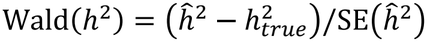. Asymptotically, Wald(*d*) and Wald(*h*^2^) are standardized, so the slope *k* of their regression equals their correlation, *k*<1. In simulations, Wald(*d*) grows approximately linearly with Wald(*h*^2^): *k* increases with true *h*^2^ but remains below 1 (Supplementary Note 7, Supplementary Table 5, Supplementary Figs 11–12). Because *k*<1, a significant Wald statistic for *d*=1 implies an even larger deviation of Wald(*h*^2^), so the test for *d*=1 flags clearly misestimated *h*^2^ but may miss moderate bias 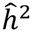. We therefore use the conventional threshold 1.96: in simulations, the test rejected *d*=1 in none of the replicates at *h*^2^≤0.05 and in 0.6% at *h*^2^=0.1.

To evaluate regularization, we used UKB LD matrices (in-sample) and 1000G LD matrices (reference) for the same 339 loci. For each locus, *z*-statistics were simulated from the in-sample LD matrix with *n*=200,000, true *h*^2^=0.025 (100 replicates), and analyzed with (*i*) the in-sample LD matrix (τ=0.999999), (*ii*) the unregularized reference LD matrix (τ=0.97, no truncation) and (*iii*) the regularized reference LD matrix (τ=0.97 with adaptive truncation guided by the Wald statistic for *d*=1) (Supplementary Note 6).

### Benchmarking on simulated individual-level data

#### Comparison with GREML and HEELS

We benchmarked FastHer using phenotypes simulated on real genotypes from 503 unrelated Europeans (1000G, phase 3 v5)^10^. After standard quality control (MAF>0.01, HWE *p*>1×10^-3^, call rate=100%), chromosome 22 was partitioned into 24 LD-independent regions using the algorithm from^14^. For simplicity, each region was randomly pruned to 600 SNPs. Phenotypes were simulated with the *--simu-qt* option (the GCTA model), varying the simulated heritability 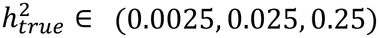 and the number of causal SNPs *m*_*causal*_ ∈ (10, *all*). For each of the 500 replicates per scenario, all methods used identical optimization settings (tolerance=1×10^-7^, maximum 5,000 iterations) and were initialized with *d*_0_=1. GREML and HEELS used a fixed initial heritability 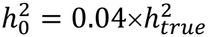, while FastHer calculate 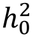 by Brent’s method at *d*=1. Here, FastHer was run as a two-parameter REML yielding the same estimates as the profile REML (Supplementary Note 1.1)

Across all scenarios, FastHer and HEELS produced practically identical estimates of *h*^2^ and its SE, in good agreement with the GREML benchmark (Supplementary Table 3; Supplementary Figs 1–6; Fig. 1a shows the estimates for the scenario 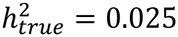, *m*_*causal*_ = *all*). These results support the accuracy of the summary-statistics approach. At low simulated heritability 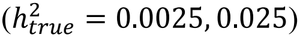, all methods were biased upward, a known phenomenon called the ‘boundary effect’: estimates falling below zero are set to zero, so the mean of the estimates exceeds the true value. The bias of FastHer and HEELS was comparable to that of GREML and negligible at 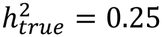 (Supplementary Table 3).

#### Comparison with SumHer

Because summary-statistics methods are designed for large GWAS, for this comparison only we enlarged the sample to *n*=5,000: the 503 genotypes were augmented with 4,497 simulated individuals drawn from a Gaussian copula with the observed LD matrix and allele frequencies (chromosome 22, 24 loci, 700 SNPs per locus, MAF>0.01). Then phenotypes were simulated under the LDAK model (--power −0.25) with all SNPs causal, 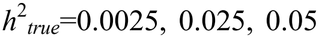, and about 100 replicates per locus. SumHer and FastHer were applied to the same genotypes; FastHer and SumHer used the same MAF-dependent weights (power −0.25) (Supplementary Note 5.1; Supplementary Fig. 9; Supplementary Table 4)

#### Analysis of d

For each replicate, we tested *d*=1 with a Wald test, using the FastHer *d* estimate and its standard error: replicates with 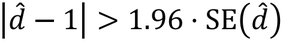 were flagged as inconsistent with the model. Results are given for all replicates and for those passing the test, the same replicates for all methods (none were excluded at any of the simulated *h*^2^ values). Calibration of standard errors was assessed as the mean SE divided by the empirical SD of the estimates.

### Runtime comparison of FastHer and HEELS

We compared the runtimes of FastHer and HEELS on 269 genes of chromosome 22 (301–10,968 SNPs) for simulated heritability 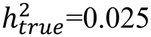 and on 266 genes with up to 9,728 SNPs for 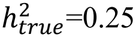, using the precomputed UKB LD matrices^15^ (*Data Availability*). *z*-statistics were simulated from a multinormal distribution with *d*=1 and all SNPs causal, with 10 replicates per gene. Computations were performed on a single Intel Xeon E5-2650 v2 2.60 GHz core. For 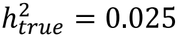, the largest gene (10,968 SNPs) took 6.4 min with FastHer and 10.7 h with HEELS (∼100-fold faster; Fig. 1c; Supplementary Note 4, Supplementary Fig. 8).

### Application to real GWAS summary statistics

We applied FastHer and HEELS to UKB-SAIGE GWAS summary statistics for five binary cardiometabolic traits: type 2 diabetes (PheCode 250.2), hyperlipidemia (PheCode 272.1), myocardial infarction (PheCode 411.2), angina pectoris (PheCode 411.3) and coronary atherosclerosis (PheCode 411.4; Supplementary Note 3). The analysis of these traits was performed for chromosome 19. We also analyzed three continuous biomarkers, alkaline phosphatase (ALP), aspartate aminotransferase (AST) and insulin-like growth factor 1 (IGF-1), for chromosome 22. For both datasets, we used gene-level UKB LD matrices^15^ (*Data Availability*). To account for variable sample sizes across SNPs, *z*-statistics were adjusted using the harmonic mean sample size.

To obtain biologically plausible estimates (low gene-level *h*^2^ and *d*≈1), we used stringent convergence criteria specific to each method. For HEELS, the tolerance was 10^−9^ for the maximum change in the variance components (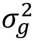 and 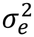) between iterations. For FastHer, the tolerance was 10^-16^ for the change in log-likelihood between iterations. Both methods were limited to 10,000 iterations. Convergence of the algorithms was monitored for every gene, and the gene-level estimates of *h*^2^ and SE(*h*^2^) from the two methods were compared (Supplementary Fig. 7).

## Data Availability

The 1000 Genomes Project reference data are publicly available at http://ftp.1000genomes.ebi.ac.uk. UK Biobank GWAS summary statistics for the cardiometabolic traits and biomarkers analysed here are accessible via the SAIGE-based PheWeb portal (https://pheweb.org/UKB-SAIGE/) and the University of Michigan data repository (https://share.sph.umich.edu/). The gene-based LD matrices were precomputed from UK Biobank data (application #59345; www.ukbiobank.ac.uk) and are publicly available at https://mga.icgbio.ru/sumfregat/ukbb/.

## Code Availability

FastHer (version 1.1.0) is freely available as an R package (https://github.com/gulsvi/FastHer) and archived on Zenodo (https://doi.org/10.5281/zenodo.24396204). For comparison, we used SumHer (https://dougspeed.com/sumher/), HEELS (https://github.com/huilisabrina/HEELS) and GCTA-GREML (https://yanglab.westlake.edu.cn/software/gcta).

## Acknowledgements

This work was supported by the budget project of the Institute of Cytology and Genetics (FWNR-2026-0023). The research was conducted using the UK Biobank Resource under application number 59345. We thank the participants and investigators of the UK Biobank and the 1000 Genomes Project for making their data publicly available.

## Author Contributions

G.R.S. conceived the method, developed the theoretical framework, implemented the software, performed the formal analysis and data curation, visualized the results, and wrote the original draft of the manuscript. Y.A.T. performed the investigation and reviewed and edited the manuscript. T.I.A. supervised the project and reviewed and edited the manuscript. All authors approved the final version.

## Ethics Declaration

This study presents a computational method and does not involve human or animal subjects. All benchmarking analyses were performed using publicly available data from the 1000 Genomes Project and the UK Biobank, which were obtained in accordance with their respective ethical regulations. Only publicly available individual-level genotypes (1000 Genomes Project) and summary-level data were used.

