## Supplementary Notes 1-7 for "FastHer: fast and accurate estimator of local heritability using summary statistics"

### Note 1 | Theoretical details

**Supplementary Table 1 | Methodological comparison of FastHer, HEELS and SumHer**

| Feature | FastHer | HEELS | SumHer |
|---|---|---|---|
| Operates directly on precomputed LD matrices | ✓ | ✓ | ✗ |
| Performs exact REML | ✓ | ✓ | ✗ |
| Estimates SE analytically using Fisher information matrix | ✓ | ✓ | ✗ |
| Imposes no MAF restriction (includes rare and ultra-rare SNPs) | ✓ | ✓ | ✗ |
| Supports user-defined SNP weights and reference LD panels | ✓ | ✗ | ✓ |
| Linear per-iteration cost | ✓ | ✗ | N/A |

✓, supported; ✗, not supported; N/A, not applicable.

##### 1.1 Initialization of the two-parameter REML

The profile REML requires no initialization: $\Phi(h^2)$ is a function of a single variable and is minimized over $[0,1)$ by a bounded scalar search. When $\Phi(h^2)$ is flat, has several local minima, or the estimate approaches the boundary of $[0,1)$, FastHer instead maximizes $l_y(h^2, d)$ jointly over both parameters, which requires initial values for $(h^2, d)$. To obtain a robust and computationally inexpensive starting point, we set $d_0 = 1$ and obtain $h_0^2$ by maximizing the one-parameter log-likelihood $l_y(h^2, d = 1)$ over $[0,1)$ with Brent's method, which needs a search interval but no initial value. The pair $(h_0^2, 1)$ then serves as the starting point for the joint optimization of $h^2$ and $d$.

##### 1.2 Standard errors

For the −2 log-likelihood (13), to calculate standard errors of $h^2$ and $d$, we derived the functions of partial derivatives and formed the Fisher information matrix. Our derivations leverage Eqs. (18)–(19) of *Methods* and standard matrix calculus rules for a matrix-valued function $X$ of a scalar $x$:

$$\frac{\partial X^{-1}}{\partial x} = -X^{-1}\frac{\partial X}{\partial x}X^{-1} \quad \text{and} \quad \frac{\partial \ln|X|}{\partial x} = \mathrm{tr}\left(X^{-1}\frac{\partial X}{\partial x}\right). \tag{20}$$

The partial first + second derivatives of $F_1 + F_2 = -2l_y$ with respect to $h^2$ and $d$ are provided in **Supplementary Table 2** (below).

**Supplementary Table 2 | First and second derivatives of $\boldsymbol{F_1 + F_2 = -2l_y}$**

| Calculus | Derivative | Expectation (true model) |
|---|---|---|
| $\dfrac{\partial(F_1+F_2)}{\partial h^2}$ | $tr(K) - \dfrac{n-m}{1-h^2} + \dfrac{n\hat{d}'(h^2)}{d}$ | 0 |
| $\dfrac{\partial(F_1+F_2)}{\partial d}$ | $\dfrac{n}{d}\left(1 - \dfrac{\hat{d}(h^2)}{d}\right)$ | 0 |
| $\dfrac{\partial^2(F_1+F_2)}{\partial(h^2)^2}$ | $-tr(K^2) - \dfrac{n-m}{(1-h^2)^2} + \dfrac{n\hat{d}''(h^2)}{d}$ | $tr(K^2) + \dfrac{n-m}{(1-h^2)^2}$ |
| $\dfrac{\partial^2(F_1+F_2)}{\partial d^2}$ | $\dfrac{n}{d^2}\left(2\dfrac{\hat{d}(h^2)}{d} - 1\right)$ | $\dfrac{n}{d^2}$ |
| $\dfrac{\partial^2(F_1+F_2)}{\partial h^2 \partial d}$ | $-\dfrac{n\,\hat{d}'(h^2)}{d^2}$ | $\dfrac{1}{d}\left(tr(K) - \dfrac{n-m}{1-h^2}\right)$ |

*Notation:* $W = \frac{nh^2}{m}U + (1-h^2)I_m$; $K = W^{-1}\left(\frac{n}{m}U - I_m\right)$; $\hat{d}(h^2)$ is given by (15), and $\hat{d}'$, $\hat{d}''$ are its first and second derivatives with respect to $h^2$:

$$\hat{d}'(h^2) = \frac{1}{n}\left[-z_u^T K W^{-1} z_u + \frac{n - z_u^T z_u}{(1-h^2)^2}\right],$$

$$\hat{d}''(h^2) = \frac{2}{n}\left[z_u^T K^2 W^{-1} z_u + \frac{n - z_u^T z_u}{(1-h^2)^3}\right].$$

Under the true model, the expectations of the first derivatives are zero, so the Fisher information matrix $I(\theta)$ for the parameter vector $\theta = (h^2, d)^T$ equals both the covariance matrix of the score and the negative expectation of the Hessian of the log-likelihood; since $-2l_y = F_1 + F_2$ ,

$$I(\theta) = -E\left[\frac{\partial^2 l_y(h^2,d)}{\partial\theta\,\partial\theta^T}\right] = \frac{1}{2}\,E\left[\frac{\partial^2(F_1+F_2)}{\partial\theta\,\partial\theta^T}\right]. \tag{21}$$

The standard errors of the estimates are obtained from the inverse of $I(\theta)$ evaluated at the REML estimates ($\hat{h}^2$, $\hat{d}$):

$$se(\hat{h}^2) = \sqrt{\left[I(\hat{\theta})^{-1}\right]_{11}}, \qquad se(\hat{d}) = \sqrt{\left[I(\hat{\theta})^{-1}\right]_{22}}. \tag{22}$$

These are Wald standard errors; they are not valid at the boundary $\hat{h}^2 = 0$, where no standard error is reported.

### Note 2 | Simulation studies: evaluating statistical accuracy (FastHer versus GREML and HEELS)

##### 2.1 Simulation design and parameters

To compare the statistical accuracy of FastHer, GREML and HEELS on simulated data, we used individual-level phenotypes generated from real genotypes from 503 unrelated European individuals (the 1000 Genomes Project phase 3 v5)[1]. After standard quality control (MAF > 0.01, HWE p > $1\times10^{-3}$, call rate = 100%), chromosome 22 was partitioned into 24 LD-independent regions (total length 35.2 Mb) with a median size of 1.33 Mb (range: 0.62–2.44 Mb) using the algorithm from[2]. Each region was randomly pruned to contain 600 SNPs. Phenotypes were simulated using GCTA's *--simu-qt* function[3] by varying two parameters: simulation heritability $h^2_{true}$ ∈ (0.0025, 0.025, 0.25) and number of causal SNPs $m_{causal}$ ∈ (10, *all*). For each of the 500 replicates per scenario, all methods used identical optimization settings (tolerance $=1\times10^{-7}$, maximum iterations = 5,000) and were initialized with $d_0$ = 1. For initial heritability values, GREML and HEELS used a fixed value $h^2_0 = 0.04\times h^2_{true}$, while FastHer (in its two-parameter version) employed its efficient initialization procedure described in **Supplementary Note 1.1.**

##### 2.2 Detailed simulation outcomes

**Supplementary Table 3 | Comparative performance of FastHer, HEELS, and GREML**

| $h^2_{true}$ | $m_{causal}$ | **Method** | **Mean** | **Bias** | **MSE×10$^{-4}$** |
|---|---|---|---|---|---|
| 0.0025 | 10 | FastHer | 0.0225 | 0.020 | 7.7 |
| | | HEELS | 0.0225 | 0.020 | 7.7 |
| | | GREML | 0.0215 | 0.019 | 7.2 |
| | all | FastHer | 0.0231 | 0.0206 | 8.0 |
| | | HEELS | 0.0231 | 0.0206 | 8.0 |
| | | GREML | 0.0221 | 0.0196 | 7.4 |
| 0.025 | 10 | FastHer | 0.0361 | 0.0111 | 7.8 |
| | | HEELS | 0.0361 | 0.0111 | 7.8 |
| | | GREML | 0.0348 | 0.0098 | 7.2 |
| | all | FastHer | 0.0357 | 0.0107 | 7.8 |
| | | HEELS | 0.0357 | 0.0107 | 7.8 |
| | | GREML | 0.0344 | 0.0094 | 7.2 |

| | | | | | |
|---|---|---|---|---|---|
| 0.25 | 10 | FastHer | 0.2570 | 0.0070 | 30.1 |
| | | HEELS | 0.2572 | 0.0072 | 30.2 |
| | | GREML | 0.2474 | -0.0026 | 27.3 |
| | all | FastHer | 0.2578 | 0.0078 | 27.0 |
| | | HEELS | 0.2580 | 0.0080 | 27.1 |
| | | GREML | 0.2506 | 0.0006 | 24.2 |

*Notation*: $h^2_{true}$: simulation heritability; $m_{causal}$: the number of causal SNPs; **Mean**: average heritability estimate; **Bias**: average deviation from true $h^2_{true}$; **MSE×10$^{-4}$**: mean squared error (multiplied by $10^{-4}$).

The results demonstrate statistical equivalence between the summary-statistics methods (FastHer, HEELS) and the individual-level-data method (GREML). FastHer produces estimates identical to those of HEELS and closely matching those of GREML, with only marginally higher bias and MSE.

#### Supplementary Figs 1–6

Supplementary Figs 1–6 present the results of performance comparison of FastHer, HEELS, and GREML in simulation studies under six different scenarios.

*Panel descriptions*: (a) Local $h^2$ estimates from FastHer plotted against those from GREML and HEELS. Red dashed line: $y = x$; (b) Analytical standard errors of $h^2$ estimates, SE($h^2$), from FastHer versus those from GREML and HEELS. Red-dashed line: $y = x$; (c) Distribution of $h^2$ estimates for all three methods. Red dashed line: true $h^2_{true}$ value. The lower and upper hinges correspond to the 1st and 3rd quartiles. Whiskers extend to the most extreme data points within 1.5×IQR from the hinge; (d) Mean squared error (MSE) of $h^2$ estimates across methods; (e) Relative efficiency of FastHer and HEELS compared to that of GREML.

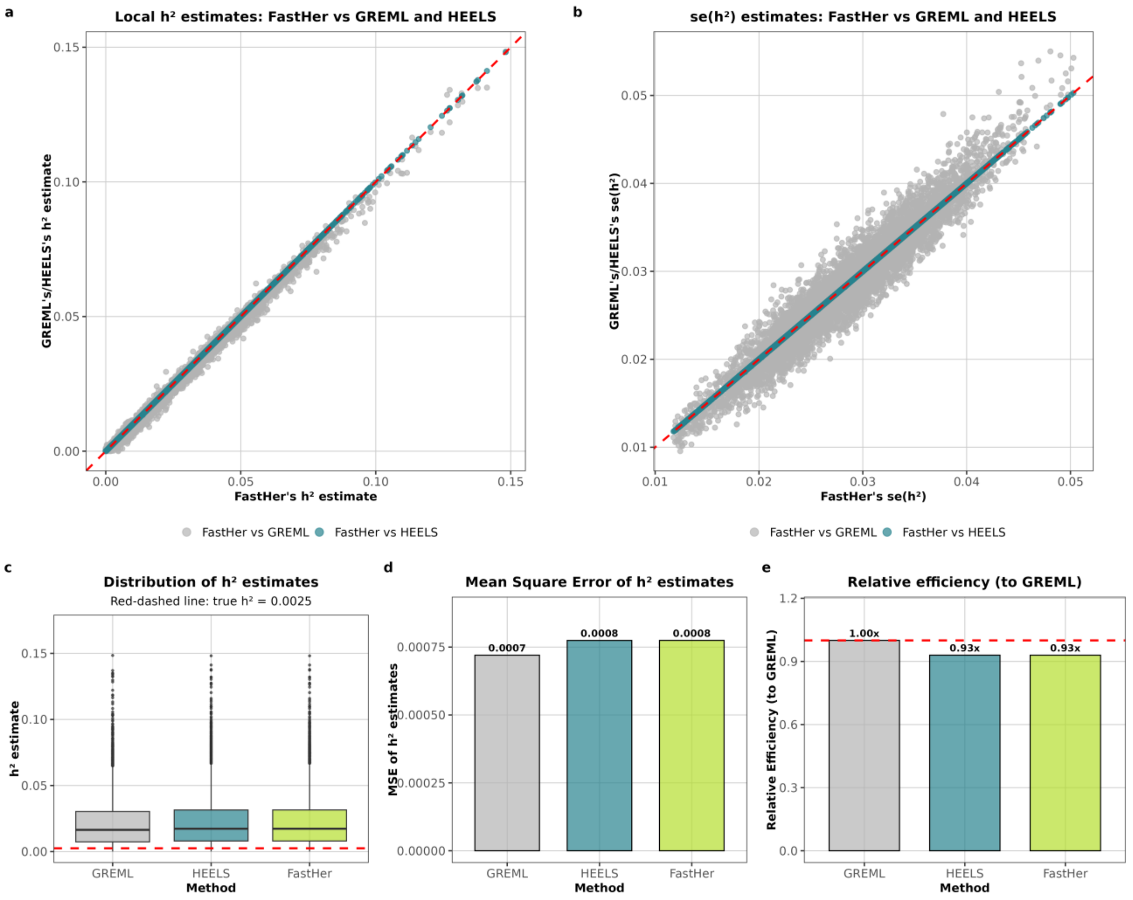


**Supplementary Figure 1. Results for scenario: $h^2_{true}$= 0.0025 with 10 SNPs as causal**

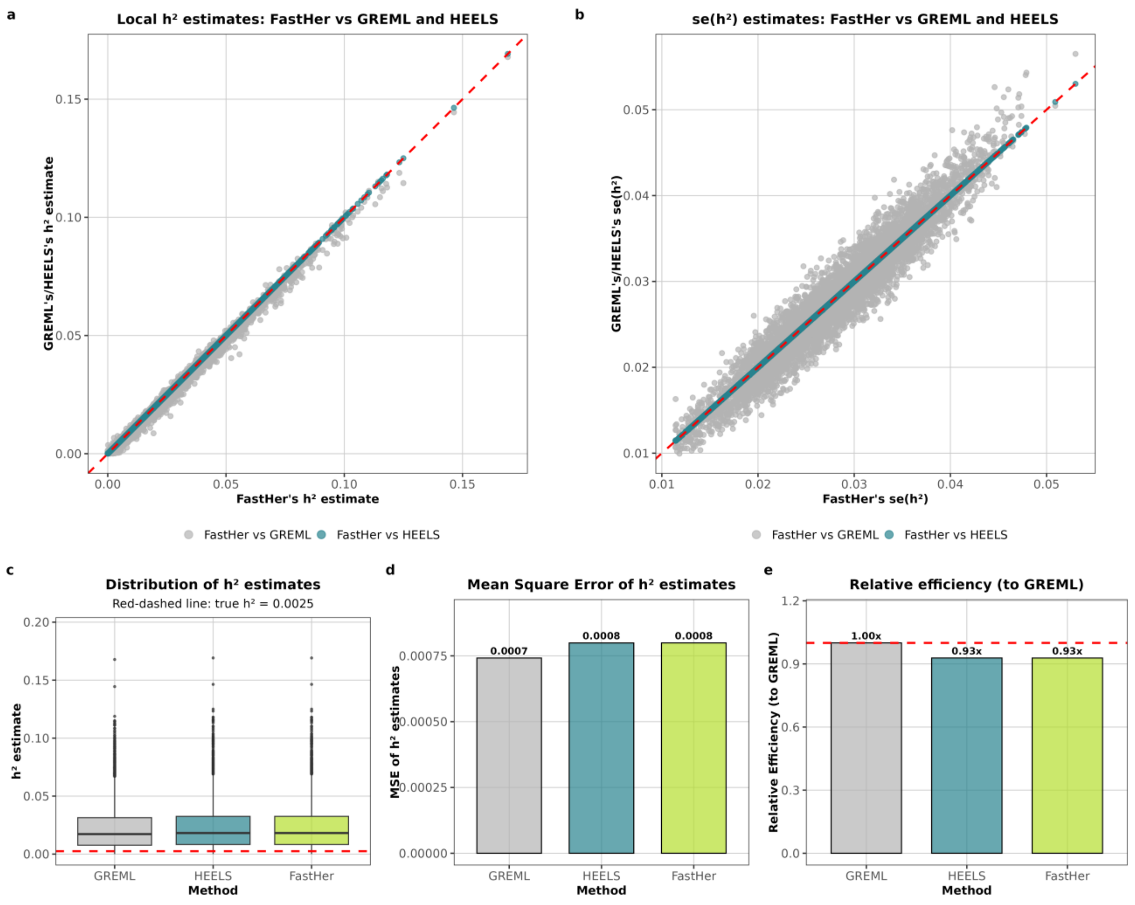


**Supplementary Figure 2. Results for scenario: $h^2_{true}$= 0.0025 with all SNPs as causal**

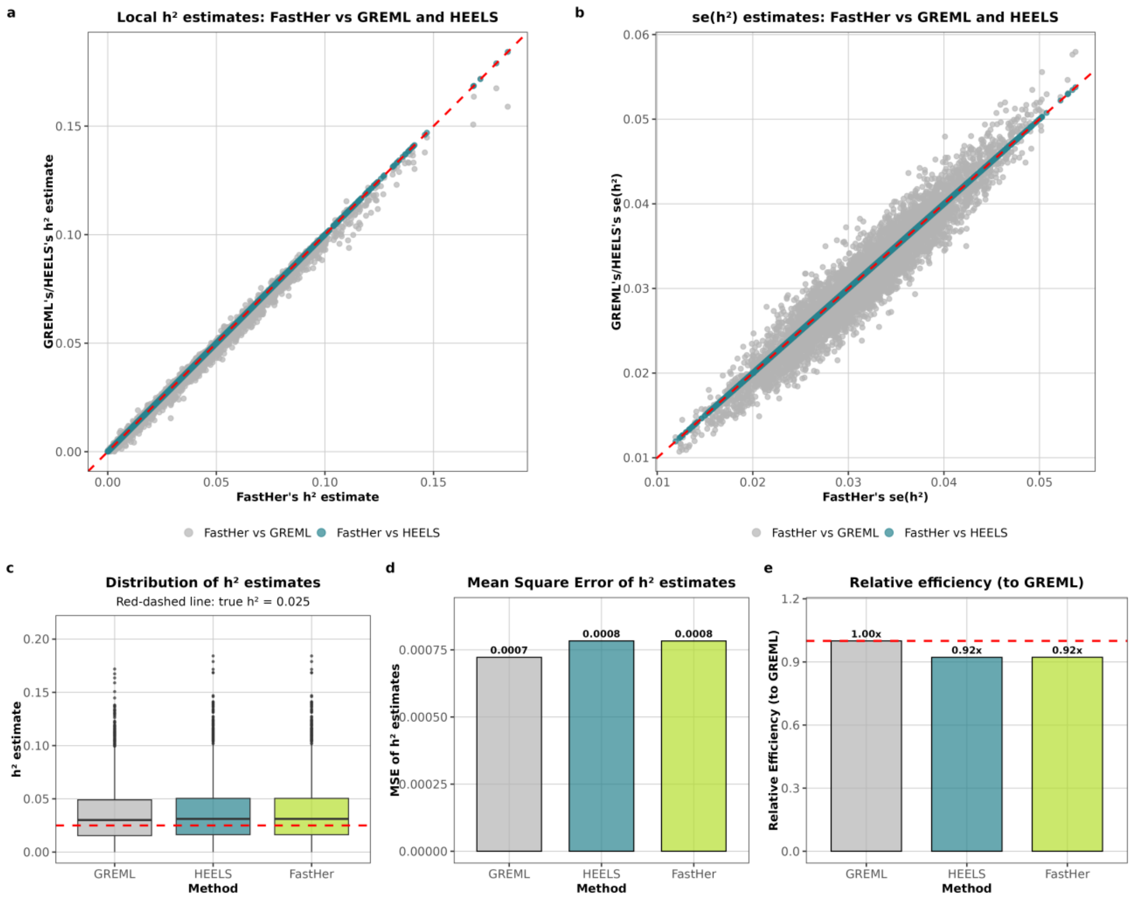


**Supplementary Figure 3. Results for scenario: $h^2_{true}$ = 0.025 with 10 SNPs as causal**

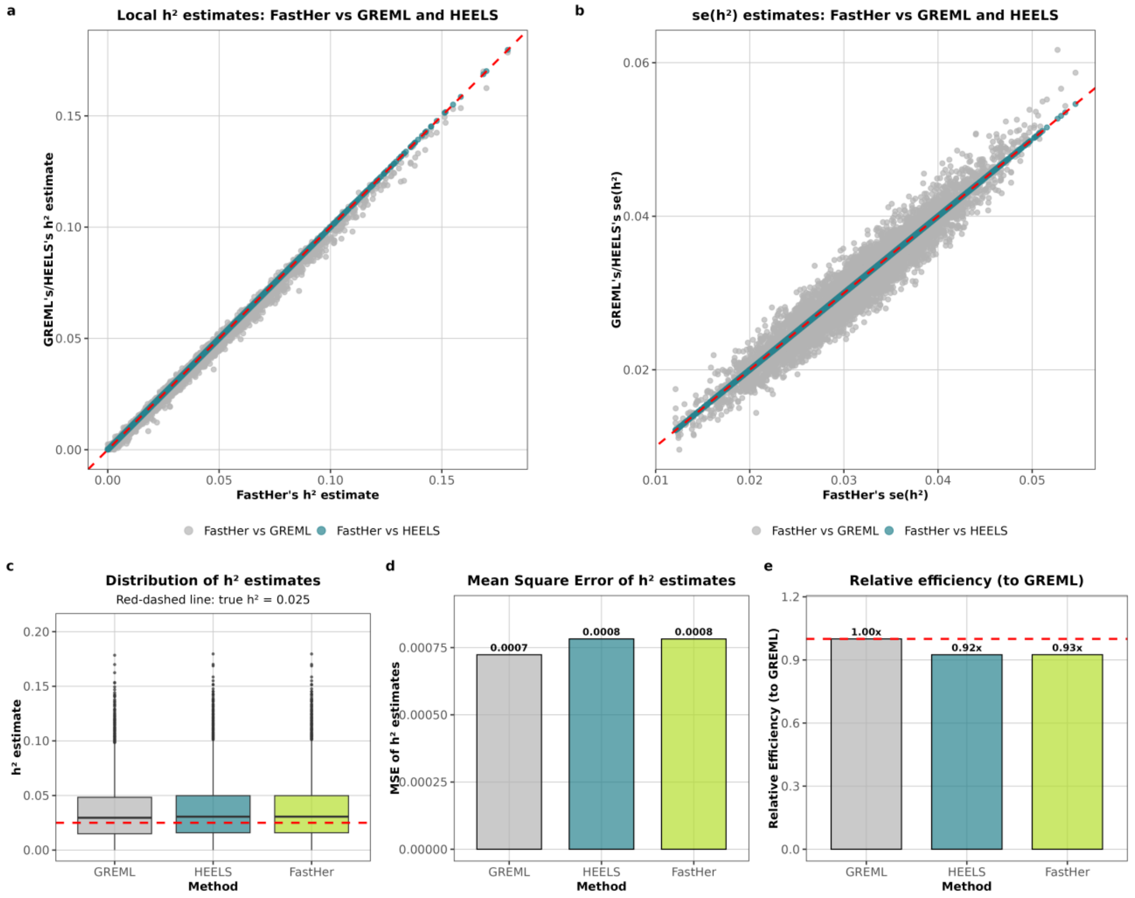


**Supplementary Figure 4. Results for scenario: $h^2_{true}$ = 0.025 with all SNPs as causal**

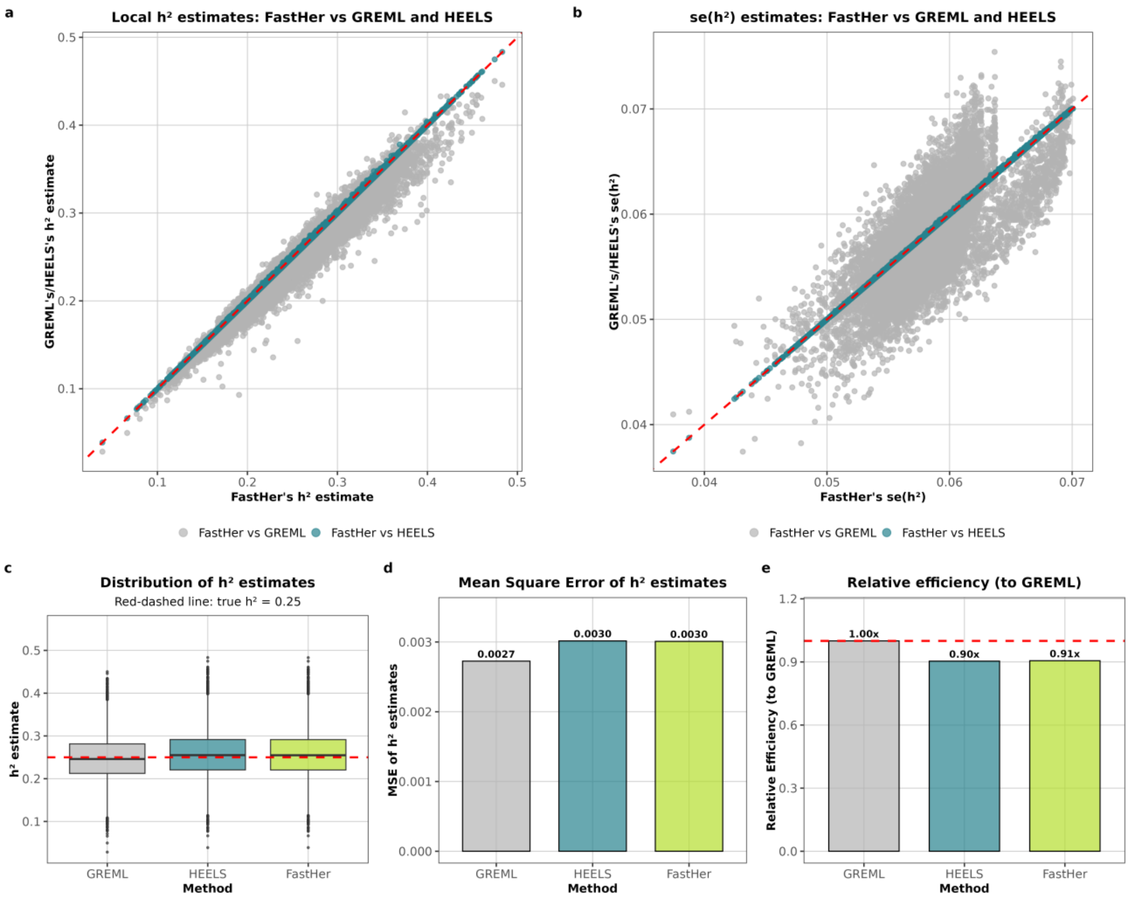


**Supplementary Figure 5. Results for scenario: $h^2_{true}$ = 0.25 with 10 SNPs as causal**

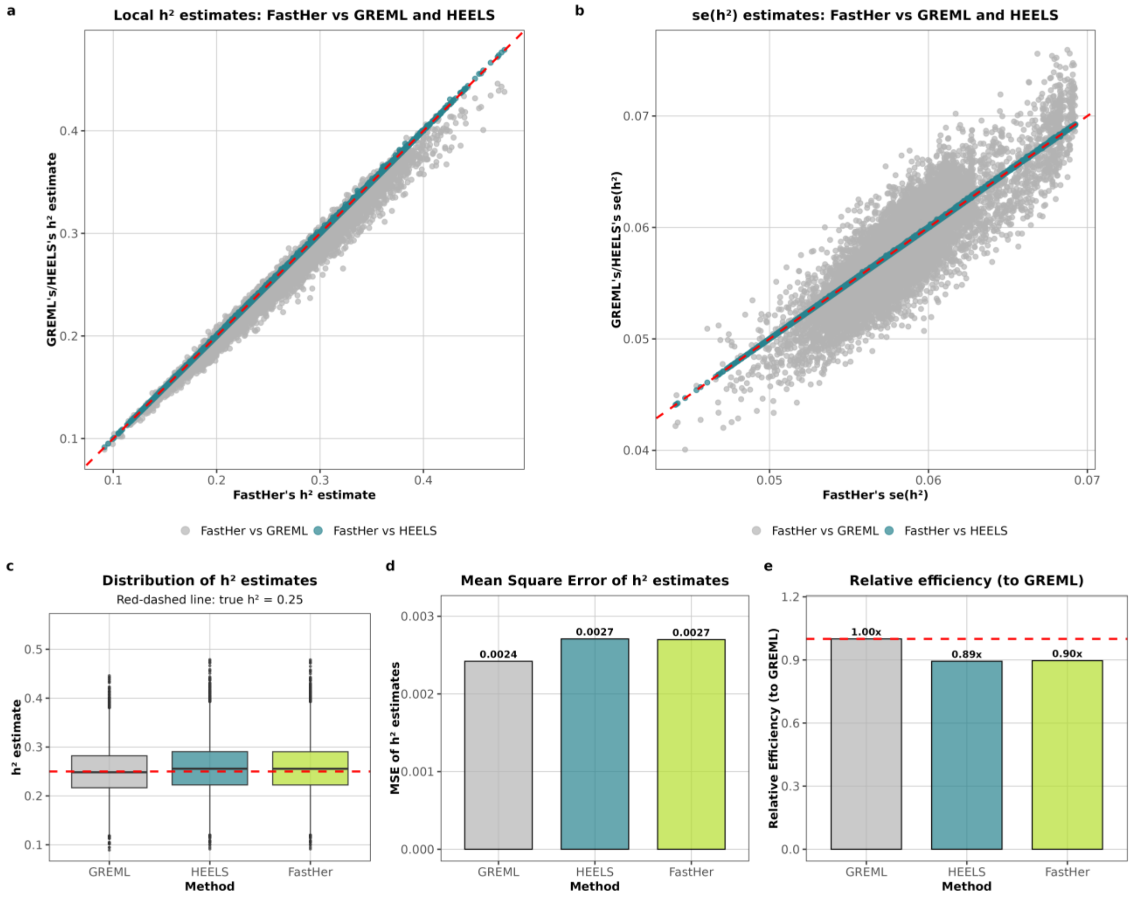


**Supplementary Figure 6. Results for scenario: $h^2_{true}$ = 0.25 with all SNPs as causal**

#### Note 3 | Comparison of FastHer and HEELS on real data

We compared FastHer and HEELS using real GWAS summary statistics from the UKB across both binary cardiometabolic diseases and continuous biomarkers.

*3.1 Binary cardiometabolic traits.* We analyzed five cardiometabolic traits from UKB using SAIGE summary statistics: type 2 diabetes (PheCode 250.2, $n$ = 407,701), hyperlipidemia (PheCode 272.1, $n$ = 408,878), myocardial infarction (PheCode 411.2, $n$ = 388,806), angina pectoris (PheCode 411.3, $n$ = 393,278), and coronary atherosclerosis (PheCode 411.4, $n$ = 397,126). The analysis used gene-based precomputed LD matrices (see *Data Availability*) from the UKB (chromosome 19).

*3.2 Continuous biomarkers.* We further validated FastHer using biomarker GWAS summary statistics from the UKB focusing on three clinically relevant continuous traits: alkaline phosphatase (ALP, Field ID 30610, $n$ = 420,633), aspartate aminotransferase (AST, Field ID 30650, $n$ = 419,034), and insulin-like growth factor 1 (IGF-1, Field ID 30710, $n$ = 418,326). The analysis utilized gene-based precomputed LD matrices (see *Data Availability*) and summary statistics for chromosome 22.

We used publicly available, precomputed gene-based LD matrices derived from 265,000 unrelated European-ancestry UKB participants. These matrices were calculated using the LDStore software[4], retaining SNPs with MAF > $1\times10^{-5}$ and imputation quality score > 0.3. Prior to analysis, z-statistics were scaled for sample size concordance using a harmonic mean approach.

To ensure biologically plausible estimates while accommodating the expected low gene-level heritability, we implemented stringent settings for convergence criteria specific to each method's algorithmic structure. HEELS used a tolerance of $1\times10^{-9}$, monitoring the maximum change in variance component estimates ($\sigma_g^2$, $\sigma_e^2$) between iterations. FastHer employed a tolerance of $1\times10^{-16}$ based on the change in log-likelihood values between iterations. Both methods were limited to a maximum of 10,000 iterations.

Applied to both binary and continuous traits, FastHer produced heritability estimates highly consistent with those from HEELS (Fig. 1b; Supplementary Fig. 7). We also observed that the runtime of HEELS depended strongly on the convergence settings and increased markedly under stringent criteria, whereas FastHer was much less affected; this observation is qualitative, as runtimes were not systematically recorded for the real-data analysis.

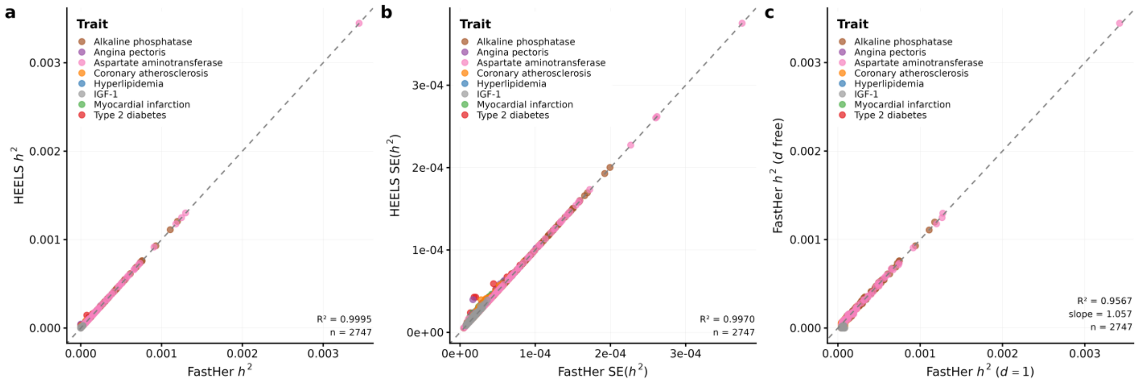


**Supplementary Figure 7 | Comparison of FastHer and HEELS methods using real GWAS summary statistics from the UK Biobank.**

In total, 2,754 gene–trait pairs were analyzed (2,402 for cardiovascular traits and 352 for biomarkers); the 2,747 pairs for which the Wald test did not reject $d$=1 are shown (Wald($\hat{d}$)≤1.96 for FastHer). The remaining seven pairs (Wald($\hat{d}$)>1.96) are not shown; when re-estimated with stringent convergence settings, all of them passed the Wald test of $d$=1. **a**, Heritability estimates ($h^2$) from FastHer ($x$ axis) versus HEELS ($y$ axis). **b**, Standard errors of $h^2$ (SE($h^2$)) from FastHer versus HEELS. **c**, Heritability estimates from FastHer with the concordance parameter $d$ estimated freely ($y$ axis) versus FastHer with $d$ fixed at 1 ($x$ axis). Each point is one gene, colored by trait (five cardiometabolic traits and three biomarkers: ALP, AST, IGF-1); dashed line, $y = x$. $R^2$ (and, for **c**, the regression slope) and the number of loci are shown in each panel.

#### Note 4 | Runtime performance of FastHer and HEELS

To compare the runtime performance of FastHer and HEELS, we used the publicly available precomputed gene-level LD matrices from the UK Biobank (application #59345), the same as in the analysis of real summary statistics (Note 3). We analyzed 269 genes on chromosome 22 containing 301–10,968 SNPs $h^2 = 0.025$ and 266 genes with up to 9,728 SNPs for $h^2 = 0.25$.

$z$-statistics were simulated from a multivariate normal distribution using $z \sim \mathcal{N}(0, UW_d)$, the GCTA model reformulated in terms of summary statistics, with $d$=1 and all SNPs causal, with 10 replicates per gene at two simulated heritability values: ${h_{sim}}^2$ = (0.025, 0.25). All computations were performed on a single Intel Xeon E5-2650 v2 2.60 GHz core.

FastHer showed a substantial runtime advantage over HEELS, which increased with the number of SNPs (**Fig. 1c**). For the largest gene (10,968 SNPs) at $h^2_{sim} = 0.025$, FastHer took 385 s (6.4 min) and HEELS 38,966 s (10.8 h), i.e., about 100 times faster; for genes with more than 10,000 SNPs (25 runs), the mean runtimes were 343 s and 31,312 s, respectively (about 91 times faster). This speed gain makes the exact analysis of loci with thousands of SNPs computationally feasible.

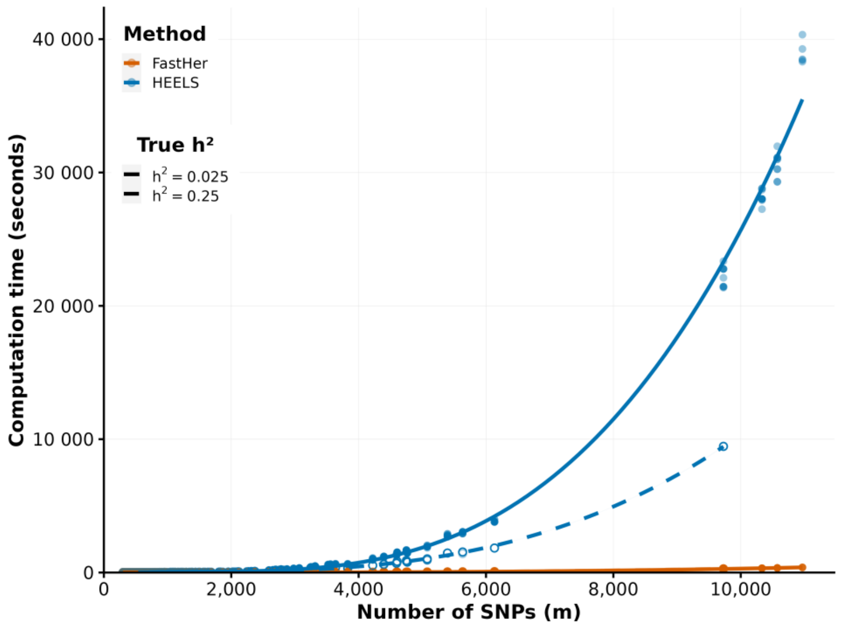


**Supplementary Figure 8 | Runtime of FastHer and HEELS versus the number of SNPs.** Computation time (seconds) for summary statistics simulated from UK Biobank gene-level LD matrices (GCTA model reformulated for summary statistics), with 10 repeats per gene. True $h^2$ = 0.025 (solid line, filled circles; 269 genes, 301–10,968 SNPs) and $h^2$ = 0.25 (dashed line, open circles; 266 genes, 301–9,728 SNPs); color denotes method (FastHer, orange; HEELS, blue). Lines are cubic polynomial fits over the measured range, with 95% confidence bands. No data for $h^2$ = 0.25 are shown for genes with more than 9,728 SNPs.

### Note 5 | Comparison of FastHer and SumHer

#### 5.1 Simulation design

*Genotypes***.** We used genotypes from the 1000 Genomes Project (phase 3, EUR, $n$ = 503) on chromosome 22. For each of the 24 LD-independent blocks, SNPs were filtered with PLINK2 *(--max-alleles 2, --maf 0.01, --geno 0.00, --hwe 1e-3, --snps-only just-acgt*), and the resulting set was thinned to 700 SNPs per locus (PLINK2 *--thin-count 700*). The 503 genotypes were then augmented to $n$ = 5,000 individuals by simulating additional genotypes from a Gaussian copula with the empirical LD matrix and allele frequencies of each locus.

*Phenotypes***.** Phenotypes were simulated under the LDAK model *(--make-phenos, --power -0.25*), with all SNPs causal *(--num-causals ALL*). Different true heritabilities were considered: $h^2$ = 0.0025, 0.025 and 0.05. Phenotypes were standardized to unit variance, and 100 replicates were generated per locus, giving 2,384, 2,400 and 2,432 replicates for $h^2$ = 0.0025, 0.025 and 0.05, respectively.

*Methods compared*. For each replicate, we applied FastHer and SumHer to the same GWAS summary statistics (z-statistics from GCTA --*fastGWA-lr*). Both methods used the same LDAK weights *(--power -0.25*; ${w_j}^2 \propto \left[MAF_j\left(1 - MAF_j\right)\right]^{0.75}$, normalized so that $mean\left(w_j^2\right) = 1$). FastHer was run with the same MAF-dependent weights; SumHer was run with --*calc-tagging* --*power -0.25* and --*sum-hers* with default settings.

*Outputs*. For each replicate, we recorded $h^2$, SE($h^2$), and (for FastHer) the concordance parameter $d$ and SE($d$). Accuracy was assessed as bias, RMSE, and relative RMSE with respect to the true $h^2$; SE calibration was assessed as mean SE divided by the empirical SD of the estimates (SE/SD; ideal value, 1; values below 1 indicate underestimated SE).

**Supplementary Table 4 | Accuracy and standard-error calibration of FastHer and SumHer, for all replicates.**

| True $h^2$ | Method | Mean | Bias | RMSE | SE/SD |
|---|---|---|---|---|---|
| 0.0025 | FastHer | 0.00294 | 0.00044 | 0.0029 | 1.10 |
| 0.0025 | SumHer | 0.00259 | 0.00009 | 0.0031 | 0.92 |
| 0.025 | FastHer | 0.0254 | 0.00042 | 0.0069 | 1.13 |
| 0.025 | SumHer | 0.0249 | -0.00012 | 0.0073 | 0.96 |
| 0.05 | FastHer | 0.0514 | 0.00141 | 0.0105 | 1.08 |
| 0.05 | SumHer | 0.0503 | 0.00030 | 0.0109 | 0.99 |

*Notation*. Bias is the mean estimate minus true $h^2$; SE/SD is the mean standard error divided by the empirical standard deviation of the estimates (values below 1 indicate underestimated SE).

FastHer and SumHer gave similar mean estimates, bias and RMSE at all three heritabilities (**Supplementary Table 4; Supplementary Fig. 9**). The ratio SE/SD ranged from 1.08 to 1.13 for FastHer and from 0.92 to 0.99 for SumHer. SumHer produced negative $h^2$ estimates in 450 of 2,384 replicates (18.9%) at $h^2$ = 0.0025, whereas FastHer constrains $h^2$ to the interval [0, 1).

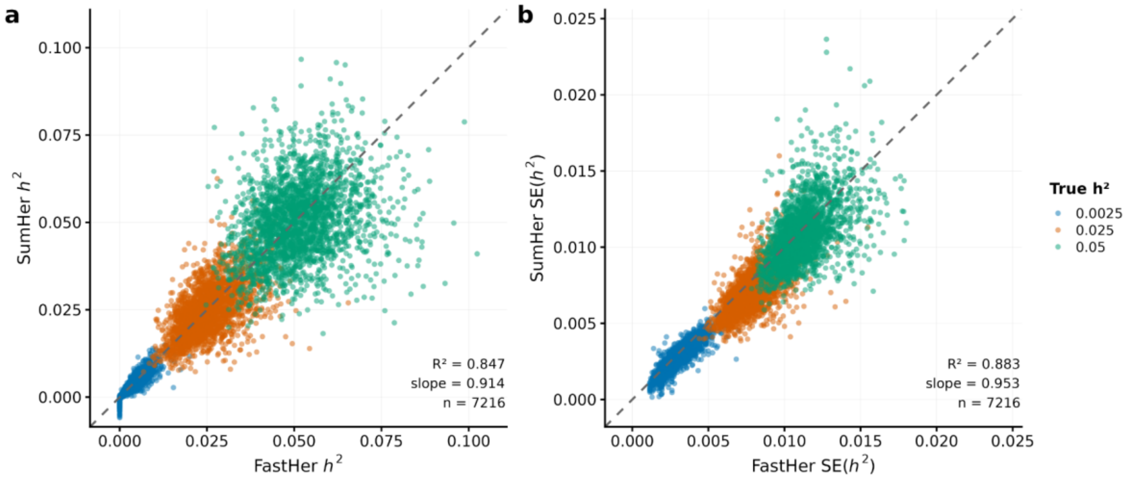


**Supplementary Figure 9 | Comparison of FastHer and SumHer across scenarios with different heritability levels.**

**a**, $h^2$ estimates from FastHer (*x* axis) versus SumHer (*y* axis) for simulated replicates, pooled across true $h^2$ = 0.0025, 0.025 and 0.05. Points are colored by true $h^2$. All replicates passed the Wald test for model-data concordance ($|\text{Wald}| \leq 1.96$ for FastHer *d*). **b**, SE($h^2$) from FastHer versus SumHer for the same replicates; color scheme as in **a**. Dashed line, $y = x$. $R^2$, regression slope and the number of replicates are shown in each panel. Both panels use identical axis scaling. FastHer and SumHer estimates were highly correlated ($R^2$=0.85 for $h^2$ and 0.88 for SE($h^2$)), with SumHer showing a slight downward bias in $h^2$ (*slope*=0.91) and close in SE($h^2$) (*slope*=0.95).

### Note 6 | Regularization of reference LD matrices

We evaluated the performance of FastHer when a reference LD matrix replaces the in-sample one. We used real UKB LD matrices (in-sample) and 1000G LD matrices (reference) for the same set of 339 loci. For each locus, we simulated summary statistics from the in-sample LD matrix, $z \sim N\left(0, \frac{n}{m} h^2 U^2 + (1 - h^2)U\right)$, with $n$=200,000 and true $h^2$ = 0.025, using 100 replicates per locus. We then analyzed the simulated $z$-statistics using three settings:

*(i)* the in-sample LD matrix (with $\tau$=0.999999);
*(ii)* the unregularized reference LD matrix (with $\tau$=0.97; no adaptive truncation);
*(iii)* the regularized reference LD matrix (with $\tau$=0.97, adaptive truncation guided by the Wald statistic for $d$=1)

Without regularization, the use of a mismatched reference LD matrix led to substantially inflated heritability estimates. With regularization, the estimates became close to those obtained using the in-sample LD matrix (**Fig. 1e**). For example, for true $h^2$=0.025, the median estimated $h^2$ was 0.0258 with the in-sample LD matrix, 0.0333 with the unregularized reference LD matrix, and 0.0257 with the regularized reference LD matrix.

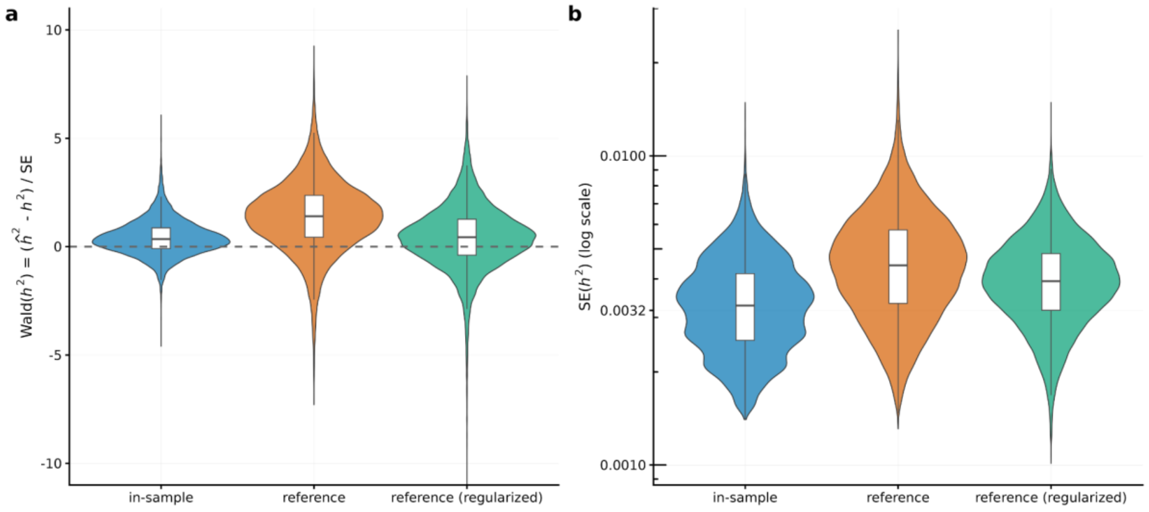


**Supplementary Figure 10 | Wald statistics and SE of $h^2$ under in-sample, unregularized reference and regularized reference LD matrices.**
Distributions of **a**, the Wald statistic for $h^2$, $\text{Wald}(h^2) = (h^2 - h^2_{true})/\text{SE}(h^2)$, and **b**, SE($h^2$) across simulated replicates (true $h^2$=0.025; 33,900 replicates per setting). The in-sample LD matrix (*blue*) yields nearly unbiased estimates (Wald median ≈0), whereas the unregularized 1000G reference LD matrix (*orange*) systematically overestimates $h^2$

(Wald median >0) and inflates standard errors. Adaptive eigenvalue truncation (*green*) corrects this bias, returning the Wald statistic to near zero, albeit with a slight increase in variance compared to the in-sample setting. Violins show the full distribution; boxplots indicate median and interquartile range. Dashed line in a indicates Wald($h^2$)=0.

### Note 7 | Concordance parameter $d$ across true heritability values

Data were simulated as in the comparison with SumHer (Supplementary Note 5.1): 24 chromosome 22 loci with 700 SNPs each, $n$ = 5,000 (503 1000G genotypes augmented with 4,497 simulated individuals), the LDAK model (--*power −0.25*) with all SNPs causal, and 100 replicates per locus (2,400 replicates per true $h^2$); true $h^2$ = 0.0025, 0.025, 0.05, 0.06, 0.07 and 0.1. For each simulated replicate, FastHer estimates the concordance parameter $d$, defined as the ratio of the model-implied marginal variance of the phenotype to its observed variance; $d$=1 indicates model-data concordance. Because $z$ and $U$ were derived from the same genotypes, $\hat{d}$ reflects the sampling uncertainty of the joint ($h^2$, $d$) estimates rather than any mismatch between summary statistics and the LD reference.

For each true $h^2$, we summarized the distribution of $\hat{d}$ (mean, median, SD, min, max, 5th and 95th percentiles), its standard error SE($\hat{d}$) (mean, median, SD), the calibration ratio $\text{mean}\left(\text{SE}(\hat{d})\right)/\text{SD}(\hat{d})$, the mean bias mean($\hat{d}$−1), and the fraction of replicates failing the Wald test for $d$=1 (|Wald| > 1.96).

**Supplementary Table 5 | Distribution of $\hat{d}$ across true heritability values, calibration and Wald test for $\hat{d}$**

| True $h^2$ | replicates | Mean $\hat{d}$ | median $\hat{d}$ | SD($\hat{d}$) | 5th pct | 95th pct |
|---|---|---|---|---|---|---|
| 0.0025 | 2400 | 0.9982 | 0.9999 | 0.0038 | 0.9903 | 1.0030 |
| 0.025 | 2400 | 1.0006 | 1.0006 | 0.0052 | 0.9921 | 1.0088 |
| 0.05 | 2400 | 1.0015 | 1.0013 | 0.0077 | 0.9894 | 1.0138 |
| 0.06 | 2400 | 1.0017 | 1.0016 | 0.0090 | 0.9874 | 1.0158 |
| 0.07 | 2400 | 1.0026 | 1.0021 | 0.0110 | 0.9856 | 1.0211 |
| 0.1 | 2400 | 1.0034 | 1.0022 | 0.0167 | 0.9780 | 1.0307 |
| **True $h^2$** | **mean SE($d$)** | **median SE($d$)** | **SD(SE($d$))** | **mean SE($\hat{d}$)/SD($\hat{d}$)** | **bias($d$)** | **Wald fail,%** |
| 0.0025 | 0.0200 | 0.0200 | 0.0001 | 5.29 | -0.0018 | 0.00 |
| 0.025 | 0.0204 | 0.0204 | 0.0003 | 3.93 | +0.0006 | 0.00 |
| 0.05 | 0.0215 | 0.0213 | 0.0008 | 2.79 | +0.0015 | 0.00 |
| 0.06 | 0.0218 | 0.0216 | 0.0009 | 2.42 | +0.0017 | 0.09 |
| 0.07 | 0.0224 | 0.0221 | 0.0012 | 2.03 | +0.0026 | 0.09 |
| 0.1 | 0.0239 | 0.0235 | 0.0018 | 1.43 | +0.0034 | 0.64 |

The results are summarized below.

1. $\hat{d}$ remained close to 1 at all true $h^2$ values (mean 0.998–1.003) – model-data concordance is preserved across heritability levels.

2. $\hat{d}$ became more variable at higher $h^2$ (SD from 0.0038 at $h^2 = 0.0025$ to 0.0167 at $h^2 = 0.1$).
3. The analytical SE($\hat{d}$) (from the Fisher information matrix) exceeded the empirical SD of $\hat{d}$ at all true $h^2$; the Wald test for $d$=1 therefore rejects less often than the nominal level, especially at low $h^2$.
4. Wald test for $d$=1: no failures at $h^2$ <0.06; 0.09% at $h^2$ = 0.06 and 0.07 and 0.64% at $h^2$ = 0.1.

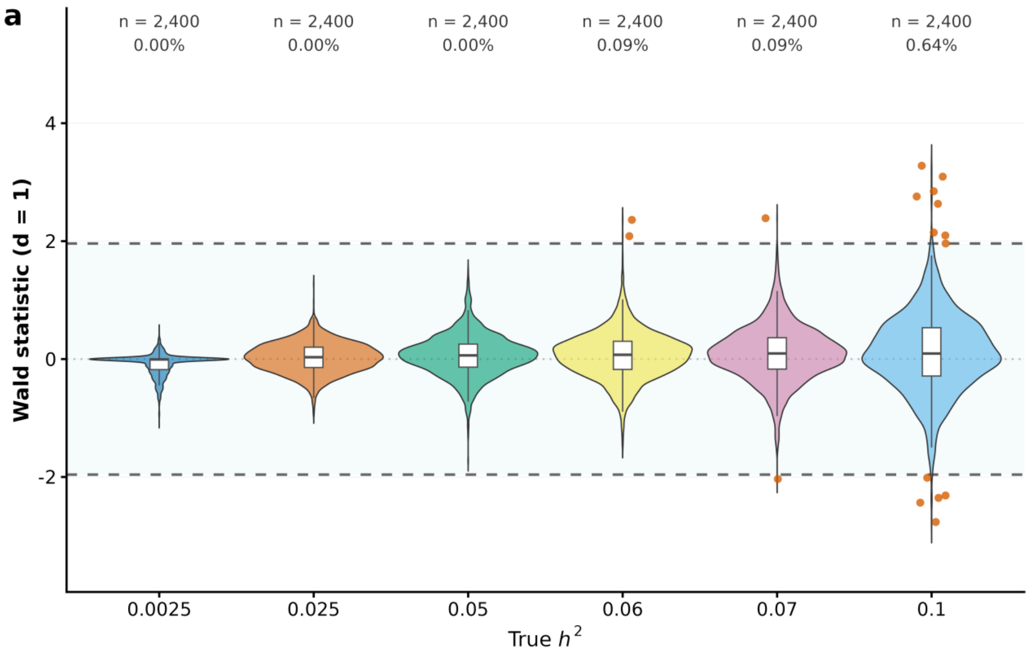


**Supplementary Figure 11 | Distribution of $\hat{d}$ (Wald statistic for $d$ = 1) across simulated replicates for true $h^2$ = 0.0025, 0.025, 0.05, 0.06, 0.07 and 0.1.**
Distribution of $\hat{d}$ (a measure of model-data concordance; $\hat{d}$ = 1 indicates full agreement) across simulated replicates for true $h^2$ = 0.0025, 0.025, 0.05, 0.06, 0.07 and 0.1. Violins show the full distribution of $\hat{d}$; white boxes show the median and interquartile range. Dashed lines mark the boundaries |Wald| = 1.96: between them, $\hat{d}$ is consistent with the model, and outside them it is not. Red points show discordant replicates (|Wald| > 1.96). Numbers above each violin give the number of replicates and the percentage of discordant replicates. The dotted line marks $\hat{d}$=1. $U$ is the LD matrix computed from the augmented genotypes ($n$=5,000), and $z$ were simulated from this matrix under the LDAK model; therefore, $\hat{d}$ reflects finite-sample fluctuations of the joint ($h^2$, $d$) estimates rather than any mismatch between $z$ and $U$.

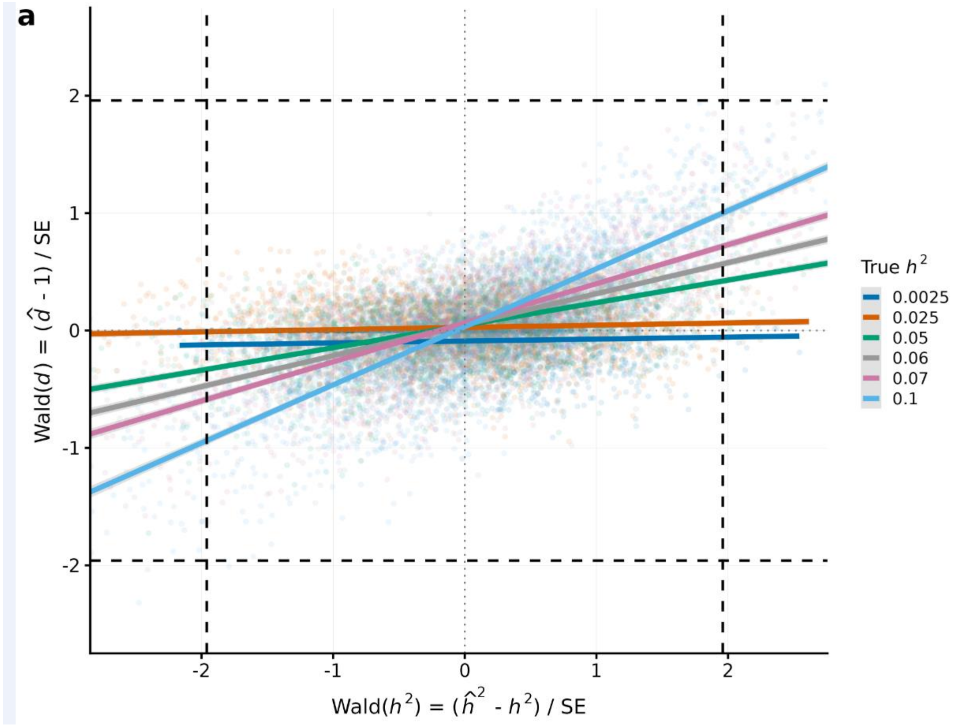


**Supplementary Figure 12 | Coupling between the Wald statistics for $h^2$ and $d$.** Wald statistic for the concordance parameter $d$=1, ($\hat{d}$–1)/SE($\hat{d}$), is plotted against the Wald statistic for heritability, ($\hat{h}^2$–$h^2$)/SE($\hat{h}^2$), for simulated replicates across true $h^2$ (color). Each point is one replicate; solid lines are linear regressions for each true $h^2$, with shaded bands indicating 95% confidence intervals. Dashed lines mark the nominal significance threshold (|Wald|=1.96; two-sided $\alpha$=0.05); dotted lines mark zero. The slope $k$ of the relationship increases with true $h^2$, from $k$≈0.02 at $h^2$=0.0025 to $k$≈0.49 at $h^2$=0.1, but remains below 1 in all scenarios (supporting the theory). Because $k$<1, a significant Wald statistic for $d$=1 implies an even larger deviation of the Wald statistic for $h^2$, and at low $h^2$ (where $k$≈0) the test produces no false positives. This allows $\hat{d}$ to serve as a practical indicator of the accuracy of $h^2$, whose true value is unknown.
